# Image memorability is predicted by discriminability and similarity in different stages of a convolutional neural network

**DOI:** 10.1101/834796

**Authors:** Griffin E. Koch, Essang Akpan, Marc N. Coutanche

## Abstract

The features of an image can be represented at multiple levels – from its low-level visual properties to high-level meaning. What drives some images to be memorable while others are forgettable? We address this question across two behavioral experiments. In the first, different layers of a convolutional neural network (CNN), which represent progressively higher levels of features, were used to select the images that would be shown to 100 participants through a form of prospective assignment. Here, the discriminability/similarity of an image with others, according to different CNN layers dictated the images presented to different groups, who made a simple indoor vs. outdoor judgment for each scene. We find that participants remember more scene images that were selected based on their low-level discriminability or high-level similarity. A second experiment replicated these results in an independent sample of fifty participants, with a different order of post-encoding tasks. Together, these experiments provide evidence that both discriminability and similarity, at different visual levels, predict image memorability.

---

When faced with the option to dine at a new restaurant, we might rely on the familiarity of a certain building, or the look of a specific logo, perhaps from a commercial we once saw. Advertising and marketing departments certainly hope this is the case, but what makes certain places and pictures more likely to be remembered than others? Determining why, when, and how images are memorable is a necessary step for modelling the relationship between visual perception and memory, and applying research, such as to select memorable health-related or educational images.

The particular images that are more likely to be remembered than others are remarkably consistent across individuals (Bainbridge, Isola, & Oliva, 2013; Bylinskii, Isola, Bainbridge, Torralba, & Oliva, 2015; Isola et al., 2014). Yet, the exact reasons that some images are more memorable than others remain to be determined. Simple visual features, such as spatial frequency, hue, and saturation, struggle to predict an image’s memorability (Bainbridge et al., 2017; Dubey et al., 2015; Isola et al., 2014), as do participants’ own subjective predictions about whether an image will be easily remembered (Isola et al., 2014). Images that are visually distinctive have been shown to be particularly memorable (Bartlett et al., 1984; Busey, 2001; Huebner & Gegenfurtner, 2012; Lukavský & Děchtěrenko, 2017), and some form of high-level content plays a role, based on the negative consequences of rearranging visual features (Lin et al., 2018) though the nature of predictive low-level and high-level content remains unclear.

The intrinsic visual properties of images play a more important role in memorability than even individual differences in observers (Bylinskii et al., 2015). An image’s properties range from low-level features such as color and edges, to high-level semantic components such as a scene’s category (Epstein & Baker, 2019). Researchers have previously quantified how visual features relate to memory with algorithms that account for the color and spatial frequencies of images (Huebner & Gegenfurtner, 2012) but such algorithms are not able to simultaneously account for higher-level properties.

Higher-level properties are thought to be central to memorability, to the extent that it has been stated that “memorability is not low-level vision” (Bainbridge et al., 2017, p. 149). Such properties have been suggested to relate to memorability even more so than visual distinctiveness (Bainbridge, 2019). It is plausible that a scene’s semantic components could be useful for predicting memory, as prior work has shown that individual objects within an image can be predictive of the overall image’s memorability (Isola et al., 2014). Little is known, however, as to how semantic properties operate within contexts, and how this might relate to memorability. Drawing on the schema and familiarity literature, we might expect that shared semantic components which are particularly important in forming a memory schema would aid memorability (van Kesteren et al., 2013). Additionally, greater semantic similarity between words (even without a visual component) tend to be better remembered (Xie et al., 2020).

Recent neuroimaging research has also highlighted the relationship between higher-level semantic features and memorability. An investigation by Bainbridge and colleagues (2017) identified that an image memorability-based similarity space is apparent in the activity patterns of higher-level visual processing regions, such as the anterior ventral stream, as well as in more traditional memory-related areas within the medial temporal lobe. High correlations between memorability and neural activity within both the visual stream and medial temporal lobe suggest that similarity is key for relating high-level properties of images to memorability. The authors noted, however, that controlling for low-level visual features in their stimulus set weakened the ability to find relationships between memorability and lower-level visual processing areas, such as early visual cortex. Thus, while these results supported claims that higher-level representations play a role in memorability, they do not necessarily rule-out a role for lower-level representations. The question therefore remains as to how lower-level *and* higher-level features (which are both present in any image) may play complementary (or even competing) roles in memorability.

Recently, rather than considering stimuli in terms of independent visual features, vision scientists have utilized convolutional neural networks (CNNs) with a hierarchical organization to characterize images (Deng et al., 2009; Kriegeskorte, 2015). These object recognition CNNs often have similarities with the human visual system (Lindsay & Miller, 2018), in which visual features are represented at increasingly high levels (Coutanche et al., 2016). Early CNN layers extract basic visual properties, which become increasingly high-level in later layers, until the final layer classifies the image (Krizhevsky et al., 2012; LeCun et al., 2015). Although primarily employed in the vision sciences, CNNs are not only applicable to perception and vision, but also to the memory field (though they are currently rarely utilized). Because CNN models allow multiple feature representations to be extracted for a single image, they make an excellent tool for measuring the impact that varied visual features may have on behavioral outcomes, such as memory performance.

In this set of experiments, we leverage the multiple layers of object recognition CNNs to determine why certain images are more memorable than others, at multiple visual stages. Uniquely, we use a CNN model to select the images that participants will view, in a prospective assignment design. Typically, research within the memory field involves collecting measures of memory performance, which are then related to others measures of interest (e.g., visual properties; Dubey, Peterson, Khosla, Yang, & Ghanem, 2015). Yet, by *retrospectively* relating memory performance to the features of items, a causal relationship between features and memory performance is not established. In contrast, prospective assignment –common in more clinical settings– provides stronger evidence of causality and helps us on the path towards explaining (Cichy & Kaiser, 2019) visual influences on memory, as opposed to reporting associations between the two. Using this prospective assignment framework allows us to better identify condition-driven behavioral differences.

Across two behavioral experiments with independent samples of participants, we test how the hierarchy of visual properties (as measured through the features at different layers within a CNN) determine an image’s memorability. We utilized prospective assignment to present participants with scene images from multiple categories that were either similar or discriminable based on CNN levels.

## Results

We investigated how image memorability is influenced by visual features at different levels.

### Experiment 1

In the first experiment, we contrasted participants’ ability to remember both discriminable and similar images of scenes, compared to foils, during a surprise recognition memory test. Participants were prospectively assigned to one of four groups in which they viewed images selected from one of four layers from the CNN. These layers correspond to different levels of a hierarchy of low-level to high-level visual features. First, we present results from analyses across the entire sample of participants, and then analyze the four groups of participants independently. On average, the odds of identifying an image as having been seen before were 18.59 times greater for the previously presented images than for matched foils (B = 2.92, *p* < .001, 95% confidence interval in odds [17.24, 20.04]). Our key question of interest was how the memory of previously presented images would differ based on which CNN layer had been used to select them. A test for polynomial effects across all 100 participants to predict memory performance for both discriminable and similar images across the four conditions (i.e., CNN layers: 1, 3, 5 or 8) revealed a significant linear interaction between image similarity (i.e., how similar an image’s features from the corresponding CNN layer are to other image’s features from the same layer) and layer condition (B = 0.33, odds = 1.39, *p* < .001, [1.16, 1.67]). This similarity x layer interaction was key to detecting the relationship, as there was no main effect of similarity when the layers were collapsed (B = 0.02, odds = 1.02, *p* = .742, [0.93, 1.11]). Neither a quadratic nor cubic function fit the data better than linear (*p*s > .197).

Each of the four conditions were then separately examined using individual regression models. Within images selected based on the earliest layer (layer 1), more similar images were less likely to be correctly recognized as old than were more discriminable images (B = −0.26, odds = 0.77, *p* = .005, [0.64, 0.93], Cohen’s *d* = 0.35). For ease of interpretability, this means that the odds of correctly recognizing an image as having been seen before were 1.30 times greater for images categorized as discriminable, than for images categorized as similar. Within the images selected based on the last layer (layer 8), the odds of correctly recognizing an image as old were 1.26 times greater for images categorized as similar, than for images categorized as discriminable (B = 0.23, *p* = .012, [1.05, 1.52], Cohen’s *d* = 0.38). No differences were observed between similar and discriminable images in the middle two layers (3 and 5) (*p*s > .581). Mean hit rates (correctly recognizing an image as old) are shown in Figure 1.

**Figure 1.**
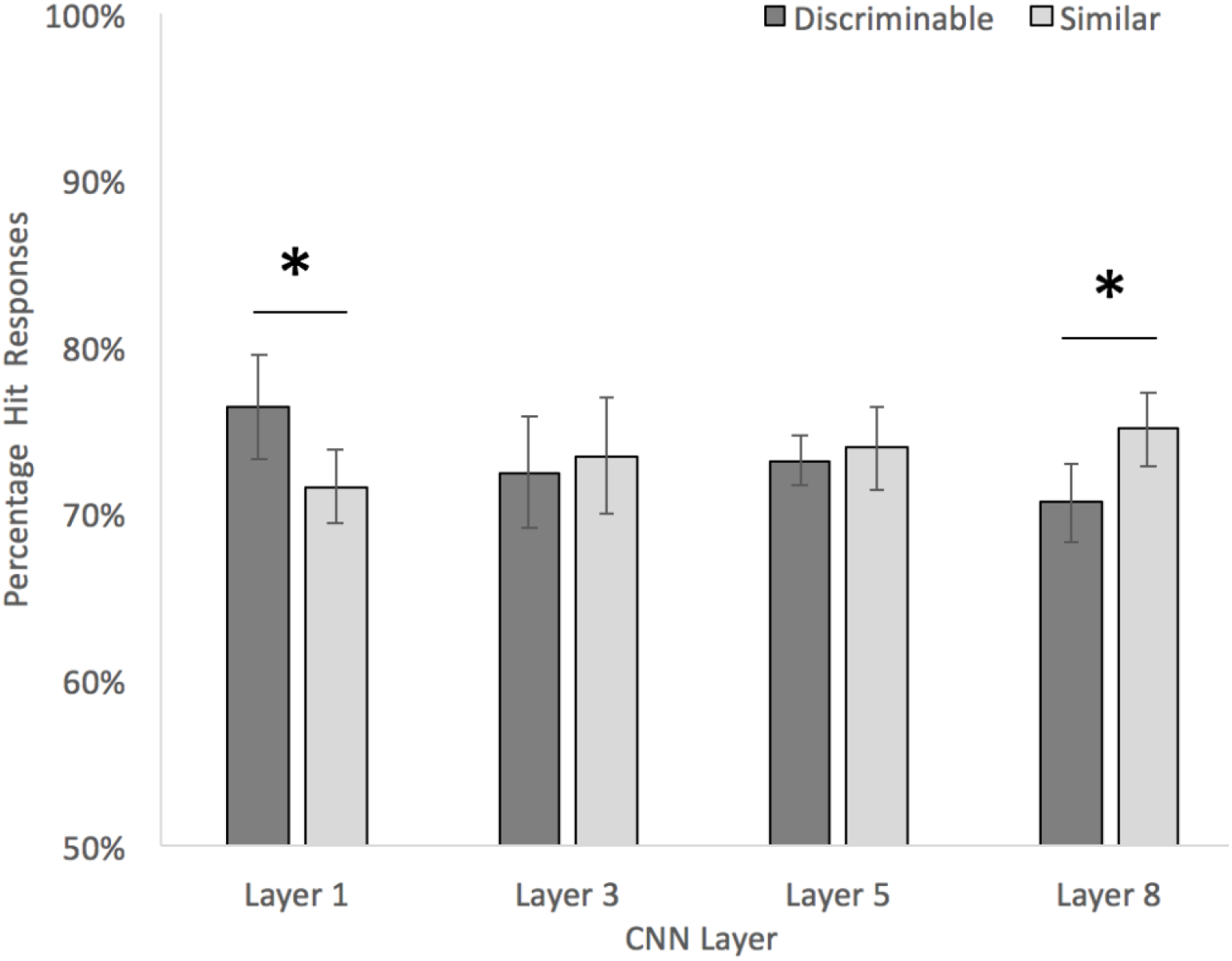
Mean hit rates for previously presented images categorized as discriminable or similar based on the four layers of the CNN for Experiment 1. Error bars reflect standard error of the mean. * represents statistical significance (*p* < .05).

We next examined whether the rate of false alarms and correct rejections of the unseen foil images differed based on the similarity/discriminability of the images for each layer. Because these foil images were not prospectively assigned in the same manner as the seen images, we assigned each foil image to have the same ‘similar’/’discriminable’ status as its corresponding seen image (from the same category). Where seen images from a category were split in being similar versus discriminable (i.e., there was no clear status to transfer to corresponding foils), the category’s foils (one category in layer 5 and four in layer 8) were removed from analysis.

Generally speaking, image similarity had opposing effects on false alarm rates as it did on hit rates. For the foils of layer 1 images, more similar images were more likely to be falsely judged as seen before, than their discriminable counterparts (B = 0.33, odds = 1.39, *p* = .004, [1.11, 1.74]). The similarity status of layer 3 foils did not affect false alarm rate (*p* = .195). In contrast to Layer 1, Layer 5 and 8 foils were more likely to be falsely judged as seen before when they were discriminable, than when they were similar (Layer 5: B = −0.49, odds = 0.61, *p* < .001, [0.48, 0.78]; Layer 8: B = −0.66, odds = 0.51, *p* < .001, [0.40, 0.66]).

We also implemented regression models with continuous (rather than similar/discriminable categorical) predictors of image similarity. For every one standard deviation *decrease* in similarity (Fisher-*Z r*-values) in layer 1, the odds of correctly recognizing an image as old were 1.14 times greater (β = −0.13, odds = 0.88, *p* = .006, [0.80, 0.96]). For layer 8, for every one standard deviation *increase* in similarity, the odds of correctly recognizing an image as old were 1.12 times greater (β = 0.11, *p* = .018, [1.02, 1.22]). No differences were observed for the middle two layers (3 and 5) (*p*s > .674).

### Experiment 2

We next asked whether these relationships between discriminability and memorability would generalize to an independent sample of participants that did not have a verbal free recall task prior to testing recognition memory (as was the case for Experiment 1). For this second experiment, we followed the same approach as in Experiment 1, but only recruited and assigned new participants to either layer 1 or layer 8, as these were the only two layers that had significant effects in the first experiment. Experiment 2 analyses therefore only investigated the lowest (layer 1) and highest (layer 8) level features, and not layers 3 or 5 (because participants were not assigned to view images from either of those layers in this second experiment). As expected based on the results from Experiment 1, participants were able to correctly recognize images that had been seen before, compared to the novel foils, with the odds of identifying an image as old being 21.98 times greater for previously presented images than for matched foils (B = 3.09, *p* < .001, [19.67, 24.57]).

As in Experiment 1, each of the two conditions were separately examined using individual regression models. Within the images based on the earliest layer (layer 1), more similar images were less likely to be correctly recognized as old than were images that were more discriminable (B = −0.71, odds = 0.49, *p* < .001, [0.41, 0.59], Cohen’s *d* = 1.33). The odds of correctly recognizing an image as old were 2.04 times greater for images categorized as discriminable, than categorized as similar. In contrast, for images selected based on the last layer (layer 8), the odds of correctly recognizing an image as old were 1.62 times greater for images categorized as similar, than as discriminable (B = 0.48, *p* < .001, [1.35, 1.94], Cohen’s *d* = 0.91). Mean hit rates (correctly recognizing an image as old) for Experiment 2 are shown in Figure 2.

**Figure 2.**
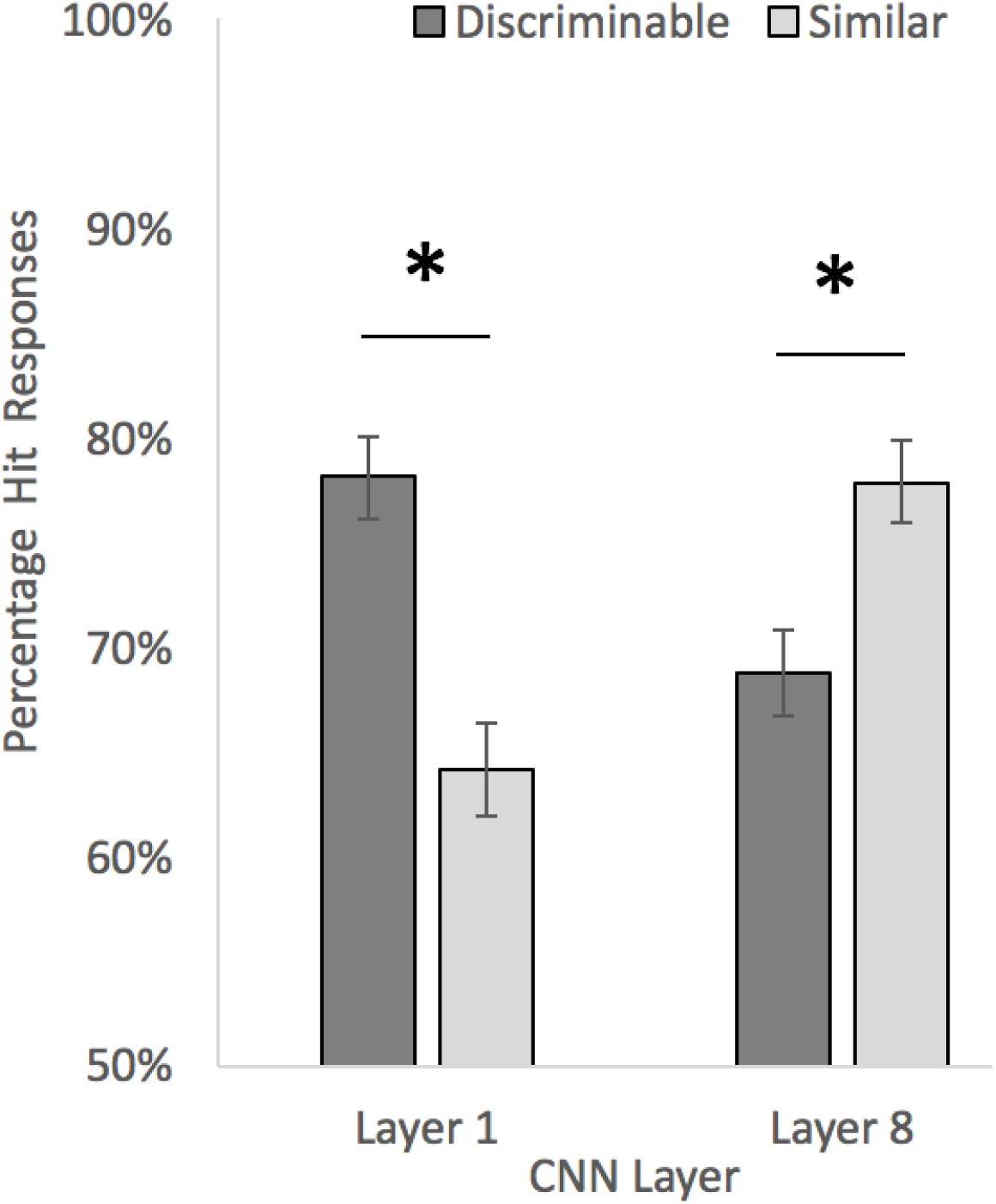
Mean hit rates for previously presented images categorized as discriminable or similar based on the four layers of the CNN for Experiment 2. Error bars reflect standard error of the mean. * represents statistical significance (*p* < .05).

As in Experiment 1, we also investigated false alarm rates. Using the same framework as described above, we again found that more discriminable layer 8 foils were more likely to be falsely judged as seen before, compared to the similar foils (B = −0.73, *p* < .001, odds = 0.48, [0.37, 0.62]. Unlike Experiment 1, more similar images in layer 1 were not more likely to be falsely judged as seen before (B = 0.12, *p* = .351, odds = 1.13, [0.88, 1.45]). These false alarm analyses only investigated the lowest (layer 1) and highest (layer 8) layers because (as discussed in the methods), participants were only prospectively assigned to images selected from these layers.

As in Experiment 1, we also implemented regression models with continuous (instead of categorical) predictors of image similarity. For every one standard deviation decrease in similarity (Fisher-*Z r*-values) based on layer 1, the odds of correctly recognizing a previously presented image as old were 1.41 times greater (β = −0.34, odds = 0.71, *p* < .001, [0.65, 0.78]). In the model using features from layer 8, for every one standard deviation increase in similarity, the odds of correctly recognizing a previously presented image as old were 1.27 times greater (β = 0.24, *p* < .001, [1.16, 1.39]).

## Discussion

We have investigated how image discriminability at multiple visual levels predicts the likelihood that an image will be remembered. In Experiment 1, we found –in a prospective assignment paradigm– that participants remembered more scene images if the images were selected based on high discriminability in low-level visual properties (earliest CNN layer), or high similarity in higher-order properties (final CNN layer). Experiment 2 replicated these results in an independent sample when the recognition test was the first test of memory after image presentation.

Our findings that similarity and discriminability can support memorability at different levels help to reconcile several seemingly conflicting findings. For instance, in some prior research, image memorability has been associated with the presence of discriminable features (Bartlett et al., 1984; Bruce et al., 1994; Lukavský & Děchtěrenko, 2017). This is in line with our evidence that discriminability within the early lowest visual level is related to better memory. Other investigations, however, have found an association between similarity (Bainbridge et al., 2017; Bainbridge & Rissman, 2018) and increased memory, which is more in line with our evidence that similarity within higher-level semantic features is related to increases in memory. Therefore, our evidence suggests that both are true – discriminability and similarity are each important predictors for whether an image will be remembered, though they operate at different stages of visual processing. Given these results, we emphasize the importance of considering the multitude of visual levels in images.

It is possible that our finding of increased memorability for images that are similar in high-level visual features could derive from links to idiosyncratic memory as suggested by Charest and colleagues (Charest et al., 2014) Such images could be the types of images that evoke the most personal episodic memories amongst people, thus resulting in increased memorability. Additionally, our finding is in line with recent neuroimaging evidence from Bainbridge and colleagues which showed that later regions within the visual ventral stream were sensitive to differences in image memorability (Bainbridge et al., 2017). These authors suggest that their findings could be in part due to feedback from memory regions within the medial temporal lobe.

The consistent results of similarity at the highest level and discriminability at the lowest level predicting memorability across both behavioral experiments highlights the robustness of these effects. Therefore, it is not likely that our findings are driven by differences in additional forms of memory testing prior to the recognition task of interest.

Our findings are in line with previous research concluding that memorability is an intrinsic property of images (Bainbridge, 2019). Both faces (Bainbridge et al., 2013) and scenes (Isola et al., 2011) have been shown to reliably elicit similar memory performances across independent groups of people. In the current study, we have shown that certain images are reliably predictive of recognition memory across two independent samples. To hone in on the images that are most memorable, we used similarity and discriminability to specifically target images at the extremes of their respective visual features (low- and high-level). These features can be measured through the use of CNNs, which extract representations of low- and high-level features after being trained on millions of images.

An aspect of the design worth highlighting is the prospective assignment performed in Experiments 1 and 2. Typically, experiments of memorability test how memory for a large set of intermixed images varies with several metrics, such as visual properties. This approach can be valuable for identifying potential underlying predictors of memorability, but by *retrospectively* relating features to memory performance, the relationship is necessarily correlational. In contrast, prospective assignment –common in clinical trials– provides stronger evidence of causality because the hypothesized dimension of interest (here, CNN layer discriminability) is used to allocate participants to different conditions in advance, and allows for greater confidence in the reason for differing outcomes (memory performance) across the groups. In addition to giving us greater confidence in the cause of differences in image memorability, prospectively assigning images to different groups also minimizes any potential interference that could otherwise occur when participants view images from multiple sets (Konkle et al., 2010a; Tulving, 1972). Future memorability research might consider prospectively partitioning presented items based on hypothesized features of interest, as opposed to the more common method of retrospectively relating memory performance to stimulus-level features in a large set of presented stimuli.

Our exploratory analyses of false alarm rates based on CNN layers provide an intriguing avenue for future research. We found that discriminability of lower-level visual features not only aids recognition of previously seen images, but also aids in successfully rejecting images that had not been seen before (though only in Experiment 1, not Experiment 2). Additionally, higher-level semantic features showed a positive relationship between similarity and successful rejection of unseen images. We note that these results should be interpreted with caution as they were not the primary focus of these studies, and therefore were not as controlled as the analyses of the previously seen images.

The use of CNNs in the field of cognitive psychology, and specifically relating to memorability, remain underutilized. For the scope of these experiments, we were interested in the underlying levels that are represented by the layers of a given CNN (in this case, AlexNet), although we note that other CNNs have shown greater similarity with human brain activity. Future research may attempt to more closely link the specific representations between CNNs and neural activity in relation to memorability, however, the specific modeling of visual processing within the human brain, while interesting, is beyond the scope of our analyses here. The current work builds on prior research which has shown the promise of adapting CNNs to identify features relevant for memorability (Khosla et al., 2015). Our work contrasts the representations at multiple individual layers, as a way to examine the contribution of each level to memorability.

## Materials and Methods

### Participants

In Experiment 1, participants were recruited until 100 contributed usable data (25 in each condition), in line with prior research investigating recognition memory for scenes (Konkle et al., 2010a, 2010b). Participants were native English speakers with normal or corrected-to-normal vision, without a learning or attention disorder, and from the University of Pittsburgh community (49 females, 51 males, mean (*M*) age = 19.6 years, standard deviation (*SD*) = 1.7 years). Four participants’ data were not analyzed after the initial encoding phase due to low task accuracy (described in more detail below), so were removed and excluded from the target of 100 participants.

For Experiment 2, participants were recruited until 50 contributed usable data (25 in each condition). Participants were recruited using the same criteria as Experiment 1, but were an independent sample (25 females, 25 males, *M* age = 19.7 years, *SD* = 1.8 years). Three participants’ data were not analyzed after the initial encoding phase due to low task accuracy (described in more detail below), so were removed and excluded from the target of 50 participants.

All participants across the two studies were native English speakers with normal or corrected-to-normal vision, and without a learning or attention disorder. The institutional review board (IRB) at the University of Pittsburgh approved all measures prior to all experiments. Participants were compensated through course credit for their participation.

### Stimuli and Materials

Stimuli for Experiments 1 and 2 were drawn from 1,000 images from the *Scenes* collection within the BOLD5000 dataset (Chang et al., 2019). Scenes ranged across semantic categories (e.g., airport, restaurant, soccer field) and included at most four images from one semantic category (e.g., four images of different soccer fields).

### Procedure

#### CNN metrics

Prior to the experiments, each scene image was submitted to the pre-trained AlexNet CNN model (Deng et al., 2009; Krizhevsky, Sutskever, & Hinton, 2012) through the MatConvNet MATLAB toolbox (http://www.vlfeat.org/matconvnet/; Vedaldi & Lenc, 2014). This CNN is one of the most well-known and accessible convolutional neural network models, and had been trained using more than 1.2 million images as part of the ImageNet object classification challenge. For each image, the CNN’s feature weights were extracted from four different layers of the CNN: earliest (convolutional layer 1), early-middle (convolutional layer 3), late-middle (convolutional layer 5), and last (fully-connected layer 8).

We focused on image discriminability (the inverse of similarity) by measuring the similarity between the presented images at each CNN layer with pairwise correlations between their sets of feature weights, giving a 1,000 × 1,000 matrix of image similarity for each layer. We Fisher-*Z* transformed the resulting correlation coefficients (*r*-values), and averaged these for each image to give a value reflecting its average similarity (or discriminability) with other images in the stimulus dataset, according to each of the CNN stages.

#### Participant assignment

For Experiment 1, participants were randomly assigned to one of four conditions. Participants in each condition were presented with images selected based on a CNN layer (layer 1, 3, 5 or 8). This subset of layers from the CNN was selected so that a diverse group of images could be presented to separate samples of participants. By spacing the layers throughout the CNN we hoped to increase the diversity of images, as representations within sequential layers will be more correlated with each other than with alternate layers.

Condition assignment dictated which set of images would be presented to a participant. The above CNN metrics of image similarity/discriminability (in each layer) were used to determine the images that were presented to each group. Each group was presented with the 50 most similar (highest *r*-values) and 50 most discriminable (lowest *r*-values) images based on its corresponding layer. See Tables 1 and 2 for a breakdown of image overlap between layers as well as similar/discriminable categorization, and Figure 3 for example images. To allow for foils from the same semantic category (defined in the stimulus dataset) to be used in the subsequent recognition task (described below), the 50 images included a maximum of two images from the same semantic category, so that a third image from the same semantic category could be used as a foil. For Experiment 2, participants were randomly assigned to one of two conditions (layer 1 or layer 8).

**Table 1.** Percentage of overlapping images

|  | Similar |  |  |  | Discriminable |  |  |  |
| --- | --- | --- | --- | --- | --- | --- | --- | --- |
|  | Layer 1 | Layer 3 | Layer 5 | Layer 8 | Layer 1 | Layer 3 | Layer 5 | Layer 8 |
| Layer 1 | 100% | 0% | 0% | 0% | 100% | 0% | 0% | 0% |
| Layer 3 | 0% | 100% | 46% | 16% | 0% | 100% | 72% | 40% |
| Layer 5 | 0% | 46% | 100% | 14% | 0% | 72% | 100% | 44% |
| Layer 8 | 0% | 16% | 14% | 100% | 0% | 40% | 44% | 100% |

Note. Values represent the number of images overlapping between layers (as a percentage of the total number of possible images). The table shows the percentage of overlapping images within those images categorized as *similar*, and within those images categorized as *discriminable* between each pair of layers.

**Table 2.**
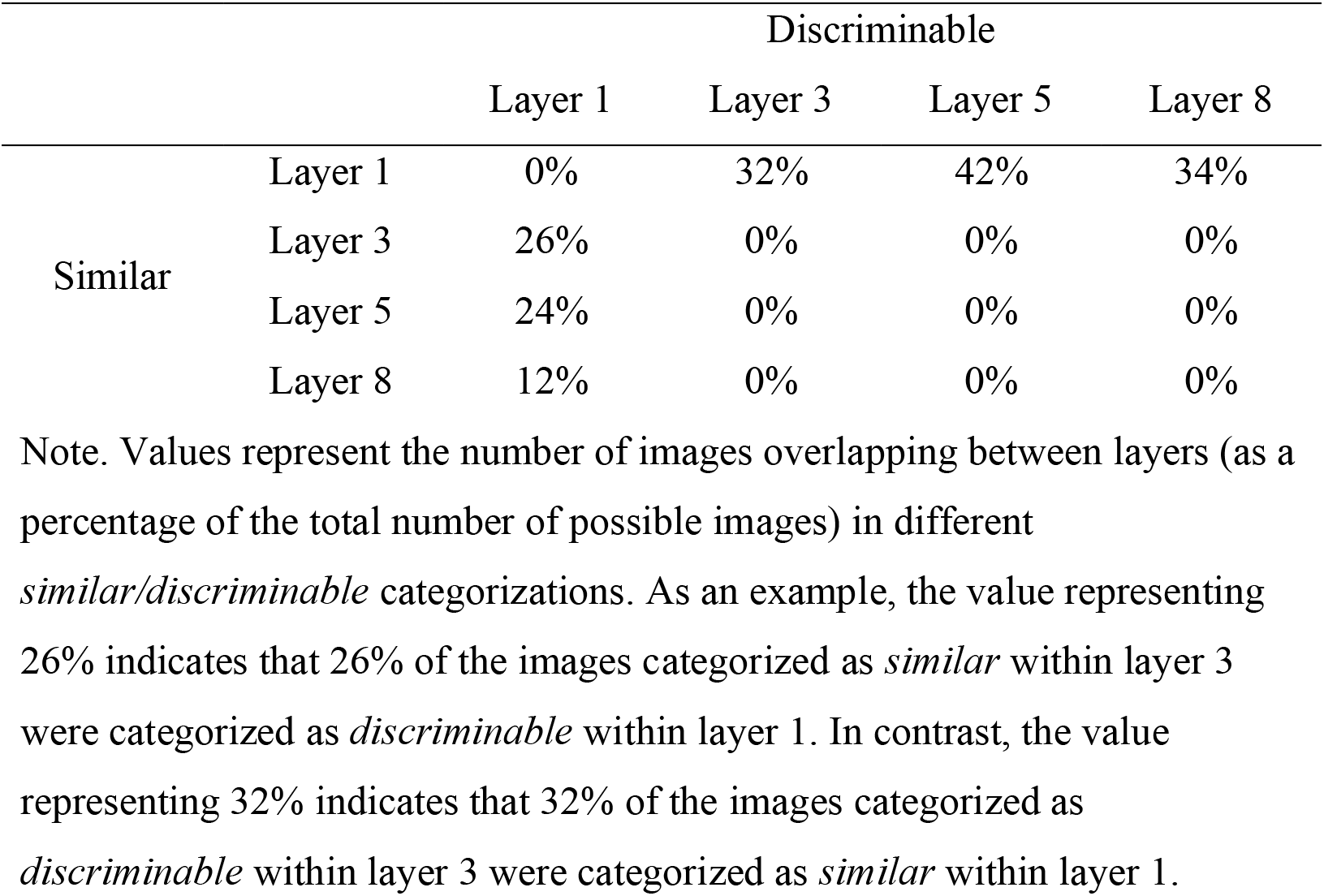
Percentage of mixed overlapping images

**Figure 3.**
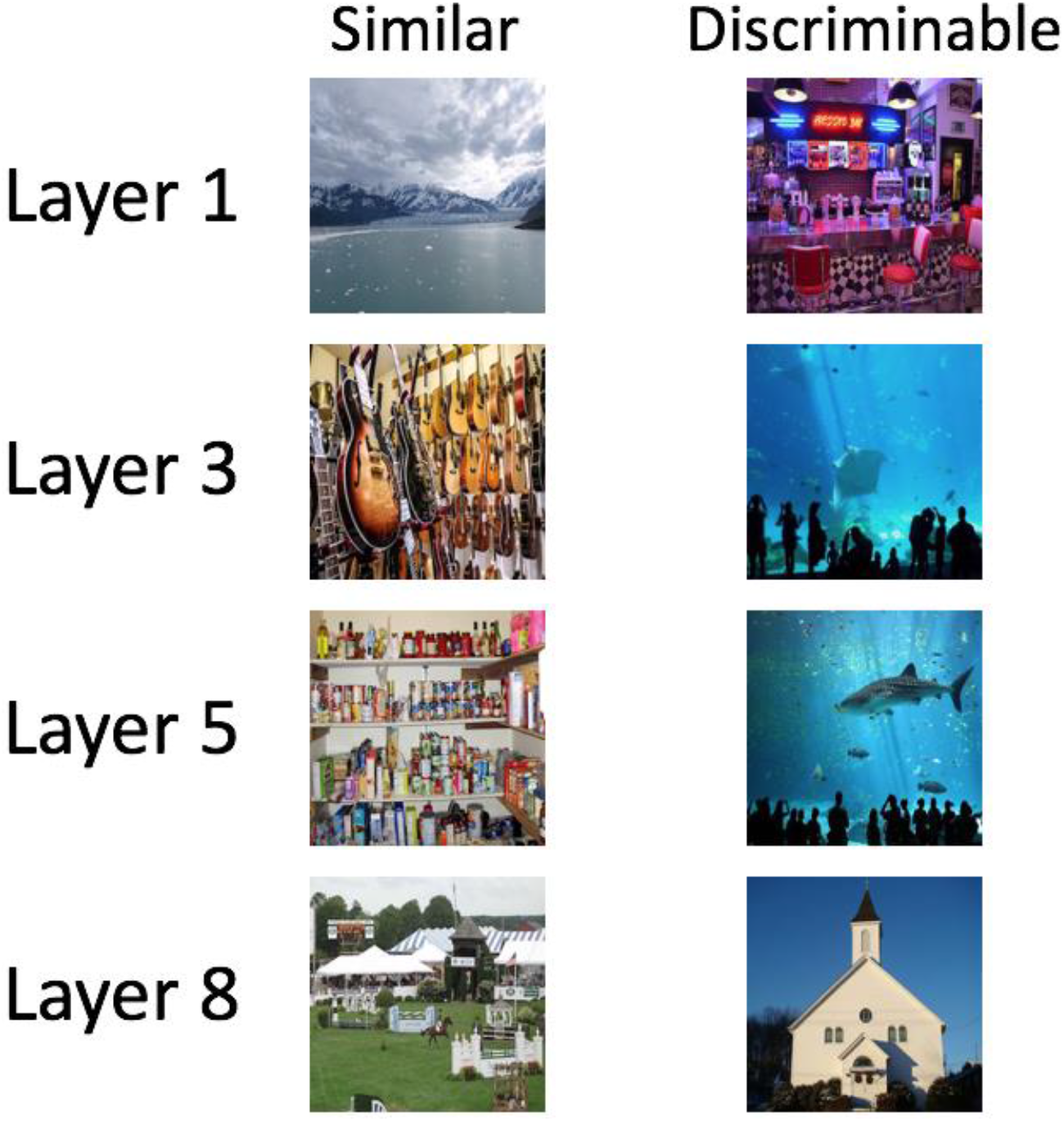
Example images matching the criteria for being selected as either “similar” or “discriminable” during Experiment 1 (layers 1, 3, 5, 8) and Experiment 2 (only layers 1 and 8). Images were categorized as either similar or discriminable from each other based on visual features extracted from a CNN. All images reproduced with permission from copyright holders. Images were resized. Attributions for images: Image of Hubbard Glacier provided via https://commons.wikimedia.org/wiki/File:Hubbard_Glacier_02.jpg; Author: James C. Space. Image of guitar store provided via https://pixabay.com/photos/guitar-store-rock-1586130/; Author: Pierre Prégardien. Image of pantry provided via https://www.flickr.com/photos/mullica/5637645692; Author: Bob. Image of Hampton Classic provided via https://commons.wikimedia.org/wiki/File:Hampton-classic1.jpg. Image of Freddy’s Bar provided via https://www.pikrepo.com/fcelk/freddy-s-bar-with-neon-lights-turned-on-and-red-stools. Image of Georgia Aquarium provided via https://www.flickr.com/photos/66087561@N08/7136330423; Author: Irish American Mom. Image of Georgia Aquarium provided via https://commons.wikimedia.org/wiki/File:Male_whale_shark_at_Georgia_Aquarium.jpg; Author: Zac Wolf. Image of church provided via https://www.publicdomainpictures.net/en/view-image.php?image=5588&picture=church; Author: Bobby Mikul.

#### Paradigm

Experiment 1 consisted of three key phases: initial encoding, free recall, and final recognition test, depicted in Figure 4. During the encoding phase, participants were presented with each of the 50 similar and 50 discriminable images of scenes from the corresponding layer (intermixed in a random order). Participants were asked to indicate whether the image was indoor or outdoor. Images remained onscreen for 4 seconds (s) regardless of a participant’s response, to allow equal encoding time across all images. A 2 s inter-trial interval followed each presented image. Upon completion of the encoding phase, participants played a game of Tetris for five minutes to prevent visual rehearsal. After Tetris, the free recall phase consisted of participants describing as many scenes as they could remember by typing as much detail as possible (not analyzed in this paper). Lastly, during a surprise recognition memory test, participants judged whether an image was old (seen previously in the experiment) or new (not seen in the experiment). The stimuli included the 100 previously seen images and 100 novel foils randomly drawn from the same semantic category as the old images (e.g., one igloo scene foil if an igloo scene was initially presented). The recognition test images were shown in a random order, and remained onscreen until participants responded (maximum 4 s). A 2 s inter-trial interval followed each image. Upon completion of this final recognition memory test, participants were debriefed about the purposes of the experiment.

**Figure 4.**
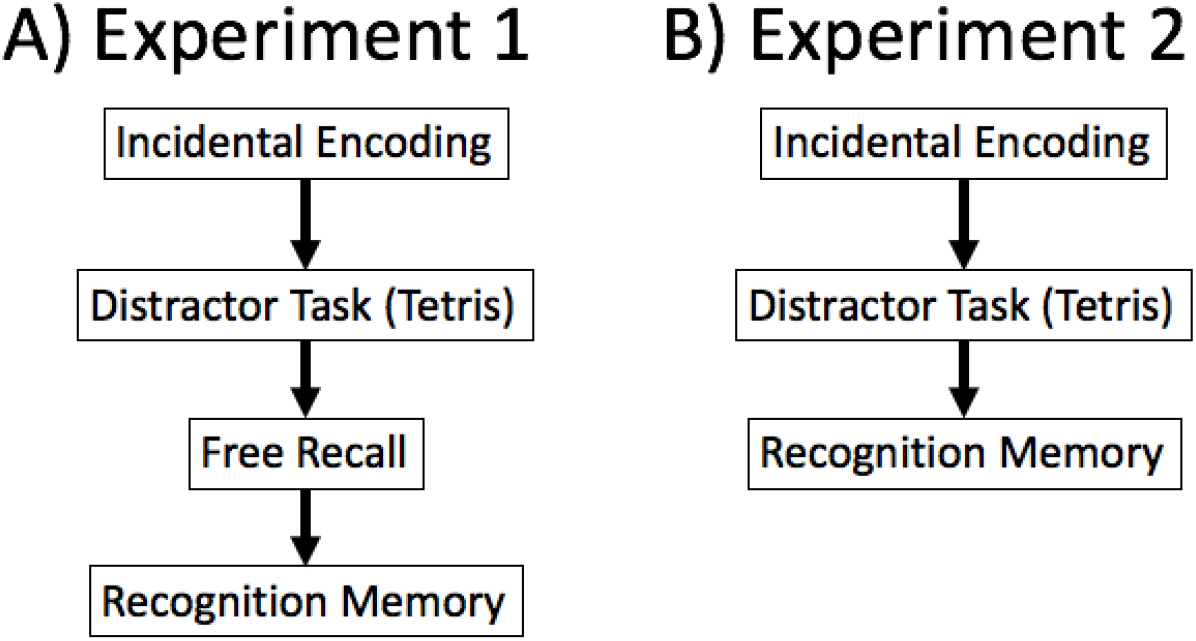
A) The experimental paradigm consisting of four tasks for Experiment 1. B) The experimental paradigm consisting of three tasks for Experiment 2.

Experiment 2 followed the same procedures as described above, except that participants did not complete a free recall phase following the game of Tetris, and instead moved directly to the surprise recognition memory test, in order to ensure that the Experiment 1 recognition memory effects were not being affected by the prior free recall test.

### Analyses

We first calculated participants’ accuracy during the initial encoding phase task (indoor vs. outdoor). Data from four participants from Experiment 1, and three from Experiment 2, were not analyzed further due to having accuracy scores that were more than two standard deviations below the mean of the full group. Behavioral results are reported based on signal detection theory implemented through logistic mixed effects regression models (Baayen et al., 2008). In each regression model, the dependent variable was the participant’s judgment as to whether or not they had previously seen the image during the encoding phase. We included fixed effects terms for image type (i.e., whether or not the image was shown during the encoding phase, and if shown, whether it was in the top 50 most similar or top 50 most discriminable for that layer), as well as the participant’s group (i.e., from which layer of the CNN the images were drawn). Additionally, a variable for participant was included as a random effect. Trials with no response were removed prior to conducting the regression models. We report unstandardized coefficient estimates (B) in logits for models with categorical predictors and standardized coefficient estimates (β) for models with continuous predictors, as well as odds and 95% confidence intervals (on the odds) as a measure of effect size.

## Acknowledgements

G.E.K. was supported by the Behavioral Brain Research Training Program from the National Institutes of Health (grant number T32GM081760). The authors thank Swathi Tata, Dan Volpone, Aarya Wadke, Grace Waldow, and Joanna Wang for their assistance in data collection. The authors also thank Scott Fraundorf for advice on statistical analyses.

